# Neurophysiological Functional Connectivity Changes during Difficult Listening in Older and Younger Adults

**DOI:** 10.64898/2026.06.06.730575

**Authors:** Vrishab Commuri, Charlie Fisher, I. M. Dushyanthi Karunathilake, Proloy Das, Stefanie E. Kuchinsky, Behtash Babadi, Jonathan Z. Simon

## Abstract

Older adults often report increased difficulty understanding speech in noisy listening environments. These difficulties are thought to arise from neurophysiological changes associated with aging, including at the level of cortex. Speech comprehension is believed to rely on coordinated activity across distributed cortical regions, mediated by the directional flow of neural signals – their functional connectivity. This connectivity can be analyzed within specific frequency bands, e.g., Delta and Theta, each thought to play a distinct role in speech and language processing. Here, we utilize the Network Localized Granger Causality (NLGC) framework applied to magnetoencephalography (MEG) data to simultaneously estimate cortical source current activities and the their directed functional connectivity. Whole-brain connectivity graphs were analyzed using graph-cluster enhancement and cluster testing. Temporal Response Functions (TRFs) were obtained by relating NLGC-derived neural current estimates to the speech stimulus presented to listeners. The band-specific networks we identify are consistent with the canonical “dual-stream” model of the auditory processing pathway. Delta-band networks primarily involve temporofrontal connections, whereas Theta-band networks showed stronger temporoparietal connectivity. These cortical networks are shown to evolve as listening conditions become increasingly adverse, incorporating frontoparietal areas associated with attention orienting and control. We find that older adults exhibit increased connectivity within areas of the ventral and dorsal auditory streams across all listening conditions, suggesting increased cortical recruitment to support speech comprehension. Although older adults exhibit enhanced time-locked responses to the speech stimulus, these responses did not significantly coincide with their connectivity patterns (and similarly for younger adults), indicating that their connectivity increases are not explained by central gain mechanisms. Together, our findings demonstrate age- and condition-dependent reorganization of speech-related cortical connectivity across neurophysiological frequency bands.

## 1 Introduction

Difficulty understanding speech, especially in complex auditory scenes, is common in aging populations. These difficulties persist even in individuals with normal audiometric thresholds (Anderson et al., 2012), suggesting neurocognitive contributions rather than arising solely from degredation at the auditory periphery.

### 1.1 Functional Organization of Cortical Auditory Processing

A central function of the brain is to integrate and interpret sensory input. Neural activity initiated at the periphery is transformed into progressively more abstract representations as the signals propagate through subcortical structures to cortical regions. This feedforward flow is referred to as “bottom-up” processing. In contrast, “top-down” processing refers to feedback signals that propagate from higher to lower stages of the hierarchy, thereby shaping auditory representations through interactions with higher-level functions such as contextual inference, attention, and sensory integration (Palmer, 1999).

Analogous to the visual system, there is evidence that auditory cortical areas are functionally divided into dual pathways or “streams” (Rauschecker and Tian, 2000; Arnott et al., 2004; Rauschecker and Scott, 2009) which convey auditory bottom-up and top-down signals. It has also been proposed that anterior ventral areas encompassing temporal and inferior frontal areas comprise a “what” pathway that is selective for lexical-semantic attributes of speech, and posterior dorsal regions consisting of temporal and parietal areas form a “where” pathway that supports motor-phonological processing (Fridriksson et al., 2016; Wang et al., 2024).

Low-frequency (1-8 Hz) neural activity from both to top-down and bottom-up processing coincide within both streams, encoding language and acoustic properties of speech (Lalor et al., 2009; Ding and Simon, 2012). Delta-band (1-4 Hz) neural activity is known to time-lock to word onsets and other slow, rhythmic aspects of speech that are closely related to its lexical and semantic attributes (Brodbeck et al., 2018a). Similarly, Theta-band (4-8 Hz) rhythms entrain to sublexical (phonemic) representations of speech that pertain to phonological components of speech (Brodbeck et al., 2018a).

In this way, structural organization and band-specific function would naturally tend to segregate, forming distinct electrophysiological networks, whose directional connections could be estimated from source-localized patterns of neural activity. A connection’s source and target each indicate a respective cortical region with a putative functional role (e.g., lower- or higher-level function), and the direction of the connection thereby can suggest a top-down or bottom-up influence. Connections can be derived from the time courses of neural activity within particular bands (Soleimani et al., 2022), so overall network topologies can be contextualized by examining aspects of the stimulus encoded in the neural activity within its nodes or within the analysis band as a whole.

### 1.2 Experimental Paradigms

#### 1.2.1 Time-Locking Paradigms

Bottom-up processing is primarily studied using paradigms that use a known stimulus and then map the elicited neural responses along the auditory hierarchy. In these approaches, neural measurements are typically averaged over many repetitions of a sound stimulus in order to capture the average evoked neural activity. The resulting response is called “time-locked” because it is temporally aligned to the timing of each stimulus event, and manifests as a sequence of positively and negatively deflected waves in the electroencephalography (EEG) or magnetoencephalography (MEG) at characteristic latencies from the stimulus onset (Luck, 2014). These M/EEG waves reflect summed electromagnetic neural activity within particular brain structures, and so by examining the timings of the observed waves, it is possible to reason about which structures generated them (for review, see Baillet, 2017). Early deflections are known to arise from subcortical and peripheral structures, whereas later deflections originate primarily from cortical regions (Luck, 2014; Skoe and Kraus, 2010; Woodman, 2010).

Time-locked responses from continuous speech stimuli can also be obtained under the Temporal Response Function (TRF) framework (Lalor et al., 2009; Ding and Simon, 2012). TRFs are linear models where the inputs are features derived from the stimulus (e.g., acoustic envelope or word timings of speech) and outputs are predicted neural response (Brodbeck and Simon, 2020; Zion Golumbic et al., 2013; Di Liberto et al., 2015).

#### 1.2.2 Source Dynamic Paradigms

In contrast, top-down processing is most typically studied by identifying how regional neural activity unfolds throughout an auditory task, independent of strict alignment to a stimulus. These approaches can be broadly termed “source dynamic” as they aim to characterize within- and inter-regional relationships by analyzing the time courses of neural source activities.

Functional Magnetic Resonance Imaging (fMRI) is the most widely used method for analyzing neural source dynamics. fMRI measures level changes in the blood hemoglobin oxygenation (the blood oxygenation level-dependent (BOLD) response) in brain tissue, instead of directly measuring electrical activity of the neurons themselves, since increased oxygenation is reflective of the increased demand of local neural activity (Amaro and Barker, 2006). In this way, fMRI can spatially localize neural responses at millimeter resolution, equipping it well to answer questions about regional recruitment during listening. Indeed, many areas outside the auditory pathway have well established functional roles relating to effort, cognition, and sensory integration (Van Den Heuvel and Pol, 2010), and their activation or interaction during an auditory task can suggest top-down involvement. However, hemodynamics vary slowly – on the order of seconds, not milliseconds – and so cannot track fast temporal changes in underlying electrophysiology. Because of this, fMRI cannot generally distinguish areas whose responses time-lock at physiological rates from those whose responses are non-time-locking but still contribute to task-specific processing (Cole et al., 2019).

### 1.3 Auditory Aging

#### 1.3.1 Time-Locking Paradigms in Aging

Several time-locking studies have reported amplitude enhancement of cortical neural responses in older listeners using evoked paradigms (Bidelman et al., 2014; McCullagh and Shinn, 2013; Tremblay et al., 2002, 2003) and the TRF framework (Brodbeck et al., 2018b; Karunathilake et al., 2023). Cortical response enhancement in older adults is thought to be at least partially caused by a central gain mechanism, by which age-related peripheral deaf-ferentation contributes to reduced inhibition and thereby produces increased cortical response amplitudes (Harris et al., 2022).

#### 1.3.2 Source Dynamic Paradigms in Aging

Source dynamic paradigms have been used to study how auditory responses in the aging brain are shaped by task demands along dimensions such as listening effort, attention mobilization, and linguistic processing. The interaction of these systems with auditory processing is typically viewed through the engagement and interaction of several functional networks in the brain. We review some of the primary networks of interest here (for a detailed overview, see Kuchinsky and Vaden, 2020).

In fMRI studies, age-related differences during speech perception often present with altered engagement of ventral and dorsal stream regions. Older adults frequently show greater activation of ventral temporofrontal regions during challenging speech listening, consistent with increased effort and reliance on lexical-semantic context (Vaden et al., 2017; Wild et al., 2012). Similarly, dorsal stream regions spanning posterior temporal, parietal, and premotor cortices may be more strongly activated in older listeners during adverse listening, potentially reflecting increased recruitment of phonological or sensorimotor integrative processing (Du et al., 2016).

The dorsal attention network comprises areas in the vicinity of the frontal eye fields (FEF) and intraparietal sulcus (IPS), and plays a role in orienting and focusing attention. Reduced intelligibility is associated with increased activation in dorsal attention networks, reflecting greater attentional mobilization during effortful listening (Peelle, 2018). These networks may also be differentially recruited in older adults during speech perception, where reduced inter-regional coordination with increasing task difficulty has been associated with poorer listening performance (Peelle et al., 2009).

### 1.4 Approach

Our work addresses two primary research questions: (1) how do cortical speech-listening networks reorganize as listening conditions become more adverse, for both younger and older listeners; and (2) how do time-locked neural responses within these networks change with increasing listening difficulty, for both younger and older listeners?

We approach these questions by means of an speech-listening experiment (Karunathilake et al., 2023) where we estimate the functional brain networks of older and younger listeners from their MEG data using the Network Localized Granger Causality (NLGC) framework (Soleimani et al., 2022). This method simultaneously obtains source-localized neural currents across the entire cortical mantle, jointly with the network of directional connections linking the sources together, which thereby avoids biases that arise in traditional two-stage approaches where source localization precedes connectivity estimation. Additionally, because NLGC is applied to MEG electrophysiological neural data obtained at high sampling rates, we obtain fMRI-like network analyses but at physiologically meaningful frequencies (viz., Delta, Theta, Alpha, Beta, Gamma). We focus here on the Delta (1-4 Hz) and Theta (4-8 Hz) bands, as these frequencies are most strongly implicated in cortical speech processing (Ding and Simon, 2012).

In order to bridge the new findings from electrophysiological studies of time-locked auditory responses with source-dynamic findings from the fMRI aging literature, we therefore characterize both network-level and response-level effects within a common experimental framework.

We hypothesize that increasing listening difficulty would produce greater recruitment of frontoparietal control networks in younger adults, reflecting improved attention mobilization under adverse conditions, as seen in fMRI. We further hypothesize that older adults would show altered network organization characterized by broader bilateral recruitment of temporofrontal and temporoparietal regions, consistent with greater engagement of ventral and dorsal auditory pathways. Finally, we expect older adults to exhibit enhanced time-locked cortical responses relative to younger adults, in line with prior reports of age-related central gain.

## 2 Methods

### 2.1 Participants

Data were collected from 18 younger (mean 20 years old) and 18 older (mean 70 years old) adults. These data were part of a previous study described in Karunathilake et al. (2023). All participants were required to have clinically normal hearing, defined as pure-tone thresholds ≤ 25 dB hearing level (HL) from 125 to 4000 Hz in at least one ear, and no more than 10 dB difference between ears at each frequency. Only participants with Montreal Cognitive Assessments (MoCA, Nasreddine et al. (2005)) scores within normal limits (≥ 26) and no history of neurological disorder were included. The study was approved by the University of Maryland’s Institutional Review Board. All participants provided written informed consent and were compensated for their time.

### 2.2 Experimental Design

Participants listened to one-minute segments from “The Legend of Sleepy Hollow” audiobook by Washington Irving. Narrations were voiced by a male or female human speaker, with both narrations level matched for equal perceptual loudness (for more details see Karunathilake et al., 2023). Segments were presented to the listener with either a single talker (speech-in-quiet) or two competing speakers. When presented with competing speech, listeners were instructed to attend to one speaker (“foreground”), and ignore the other speaker (“background”).

The speech segments were presented at three levels of listening difficulty: speech in quiet (SIQ), 0 dB SNR (level-matched competing speech), and −6 dB SNR (foreground speaker 6 dB quieter than background). A fourth condition with speech-like babble background was not analyzed in this study. Participants were instructed to minimize body movements and to attend only to the foreground speaker. During the task, listeners fixated on a projection of a cartoon male or female face which indicated which speaker to attend.

Stimuli were delivered to participants with E-A-RTONE 3 A tubes (impedance 50 Ω), which strongly attenuated frequencies above 4 kHz, and was outfitted with E-A-RLINK (Etymotic Research, Elk Grove Village, United States) disposable earbuds. Stimulus sound level was calibrated to approximately 70 dBA sound pressure level (SPL) using 500 Hz tones, and was equalized to obtain an approximately flat response from 40 Hz to 4 kHz.

The experiment was divided into two blocks. Each block began with a brief resting state scan where participants were instructed to remain still for 90 seconds. This was followed by the two competing talker (0 dB and −6 dB) conditions. The order of the competing-talker conditions was counterbalanced across participants based on foreground speaker sex and competing speech SNR (a 2 × 2 design). In the competing talker speech conditions, each stimulus was presented sequentially three times. Each block concluded with the foreground and background speech stimuli in that block presented alone as single talker speech without repetition.

### 2.3 Data Recording

Prior to MEG data recording, each participant’s head shape was digitized using Polhemus 3SPACE FASTRAK digitizer. The head position was tracked at the start and end of the experiment with 5 marker coils attached to their heads.

MEG data were recorded in a dimly lit, magnetically shielded room with 160 axial gradiometer whole head MEG system (KIT, Kanazawa, Japan) at the Maryland Neuroimaging Center. The MEG data were sampled at 2 kHz, low pass filtered at 200 Hz and notch filtered at 60 Hz. Participants laid supine during the MEG experiment while their head was in the helmet and as close as possible to the sensors.

### 2.4 Data Preprocessing

Subjects’ head position data and head shape digitization were coregistered with the Freesurfer ‘fsaverage’ brain (Fischl, 2012), which forms the cortical surface on which neural current sources are estimated. The fsaverage brain is a template brain derived from the average cortical geometry of 20 older adults, 10 middle-aged adults, and 10 younger adults. For each subject, a transformation matrix was computed using utilities provided by the MNE-python package (version 1.10.1) (Gramfort et al., 2013). The transformation encoded the rotation and translation operations needed to align the fsaverage template brain with the participant’s head position in the MEG scanner. The fsaverage brain was scaled so that it best fit the subject’s fiducials and digitization in its length, width, and height.

MNE-python was also used to perform MEG signal preprocessing. First, known noisy sensors were removed from the data, and temporal signal space separation (tSSS, Taulu and Simola (2006)) was used to regress out artifacts caused by signals originating from outside the head. The data were then filtered between 1 Hz and 45 Hz using a causal minimum phase FIR filter. Independent Component Analysis (ICA, Hyvärinen and Oja (2000)) was applied to the filtered data, and enough components were generated to account for 99% of the variance in the data. Some of these components captured cardiac and muscle artifacts which were visually identified and removed from the data (typically resulting in the exclusion of two to four components). The resulting data were further filtered to the desired frequency bands (1-4 Hz and 4-8 Hz) using causal minimum phase FIR filters, and the sampling rate was reduced to 25 Hz. For each minute-long trial, the first 5 seconds of data were discarded to account for transient responses at the beginning of the task, and the remaining 55 seconds were used in the analysis, resulting in 20 55-second trials per subject.

### 2.5 NLGC Model Fitting

Network-Localized Granger Causality (NLGC) is a state-space modeling framework that treats neural sources as latent (internal, or unobservable) states that are indirectly observed via sensor measurements from outside the head (Soleimani et al., 2022). In NLGC, latent neural sources are governed by steady-state dynamics, meaning that at any given time, a source is a function of itself and possibly other weighted contributions from other sources at previous time steps. These interconnecting influences can be thought of as a fixed network formed between the neural sources.

NLGC estimates latent neural sources using a sparse vector auto-regressive (VAR) model. The VAR model order was identified by fitting the entire data set using several candidate orders and comparing the models’ Akaike Information Criterion (AIC), a measure of model quality that jointly penalizes high model complexity and poor model fit. Nearly all subjects presented an optimal model order of 2 at a sampling rate of 25 Hz, determined based on the minimum model AIC. This order was fixed for all models to enable cross-subject comparison.

NLGC uses a lead-field summarization procedure whereby the dominant leadfield characteristics of a high-resolution source space are approximated in a lower-resolution source space. Within each cortical patch in the low-resolution ico-1 source space, leadfields for all current dipoles in high-resolution ico-4 source space were captured by principal component analysis (PCA). Our analysis retained the top four PCA components, called eigenmodes, which were stored as time-varying vectors centered within each ico-1 cortical patch. Four eigenmodes were chosen to trade off the quality of the lead field summarization with the ill-posed nature of estimating many neural sources from a limited number of MEG sensors.

### 2.6 Source Dynamics: Whole Brain Connectivity

NLGC-based connectivity maps were obtained for each subject, for each trial, and for each frequency band, resulting in twenty connectivity maps per subject per band. Each connectivity map is a matrix in which each element denotes a directed connection between cortical patch pairs.

#### 2.6.1 Normalization and Aggregation

NLGC link counts were first rate-normalized, meaning that each connectivity map was divided by its total number of links. The purpose of this operation was firstly to diminish the influences of trials which produced a spuriously high number of links, indicating that the model overfit the data and failed to adequately condense the estimated neural activity into a sparse network. A second effect of this operation was to transform connectivity counts, which vary by subject or measurement SNR, into connectivity densities that can be compared across subjects and conditions. At this stage, trials with fewer than 25 links were discarded so as not to skew the connectivity density comparisons towards overly-sparse models that underfit the data.

Connectivity is estimated for each subject in ‘ico-1’ surface source space, meaning that each link connects two of the 84 patches that tile the cortical surface. The imposition of sparsity condenses neural activity that may span multiple patches into a point source at a single patch. This results in slight misregistrations when comparing sources across trials and individuals. The need for exact localization was relaxed in two ways. First, a finite impulse response (FIR) Bartlett window with a radius of 3 cm was convolved with the source and target nodes of each link. The purpose of this was to retain the source and target centerpoint while allowing some controlled signal spreading into spatially adjacent patches. An FIR window was used to prevent leakage into distant patches.

The smoothed 84-patch ‘ico-1’ source space was then downsampled to 42 patches to further increase tolerance to small mislocalizations. To do this, a spatial adjacency graph was first constructed using the euclidean distance between cortical patch centers. Downsampling by greedily grouping together neighboring vertices will almost always result in one or more singleton vertices when all of the neighbors of a given vertex are claimed. This is a classic problem in graph theory called “maximum matching” and its solution is given by Edmonds’ Blossom Algorithm which we employed here. The result was a single downsampling transform that was applied to each native 84 × 84 connectivity map to reduce its dimension to 42 × 42, where each node in the downsampled map corresponds to exactly two nearest-neighbor nodes in the native matrix.

#### 2.6.2 Whole-Brain Connectivity Regressions: Model Definitions

Our whole-brain connectivity modeling approach draws inspiration from fMRI analysis methods that first fit simple linear models that relate regressors of interest to single observation units (e.g., hemodynamic responses within single voxels) and then test topologically-clustered effects (e.g., across voxels) for significance.

Hurdle lognormal mixed-effects regressions were fit to connectivity densities in each cell of the connectivity maps after normalization and aggregation, with one model per cell. Lognormal distributions commonly arise in biological models in which a number of nonzero influences contribute multiplicatively to produce a measured response. Because our NLGC link distributions are sparse by design, the connectivity densities have many zero-valued measurements. The hurdle component models these zero values separately (as a point-mass), distinct from the positive distribution that is fit to nonzero measurements. Generally, mixed-effects models enable investigation of how covariates of interest, e.g., age or listening condition, relate to observed connectivity while providing a framework to account for within-subject variability and a dearth of observations inherent to sparse modeling. The regression models fit to connection densities from region *i* to region *j* are of the form:

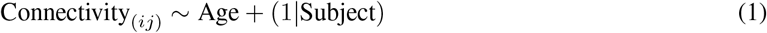

or

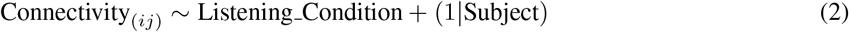

where (1|Subject) denotes a per-subject random intercept, allowing some subjects to have higher baseline levels of connectivity than others while minimally influencing the fixed effect. Note that self-connections (*i* = *j*) are permitted due to aggregation over cortical patches.

Regressions were fit to cells that contained at least 50% nonzero values. This established a support containing enough data to obtain a reasonable model fit. Posterior predictions from each model were compared to the count data to assert goodness of fit.

Models were implemented using Python’s PyMC library (Abril-Pla et al., 2023) using Monte-Carlo sampling with 4000 samples using the NUTS sampler with four chains after discarding the first 1000 samples for burn-in. Computation was performed over several days on a distributed compute cluster.

#### 2.6.3 Whole-Brain Connectivity Regressions: Null Construction

After fitting the regression model to the observed data, we characterized the influence of age or condition by comparison to a null distribution, which involves isolating the influence of the covariate of interest while not altering other parts of the model. In the language of Structural Causal Modeling, such a modification is called an “intervention”, or a *do*-operation, on the model graph (Pearl, 2009). The model graph is a representation of the hierarchical structure of the regression that links dependencies among parameters. The *do*-operation can be viewed as deleting corresponding edges in the graph, thereby altering influences of the covariate while preserving all other dependencies in the model.

An example of such an intervention on our age model is shown in Figure 1, with the corresponding generative model defined as:

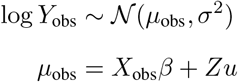

**Figure 1:**
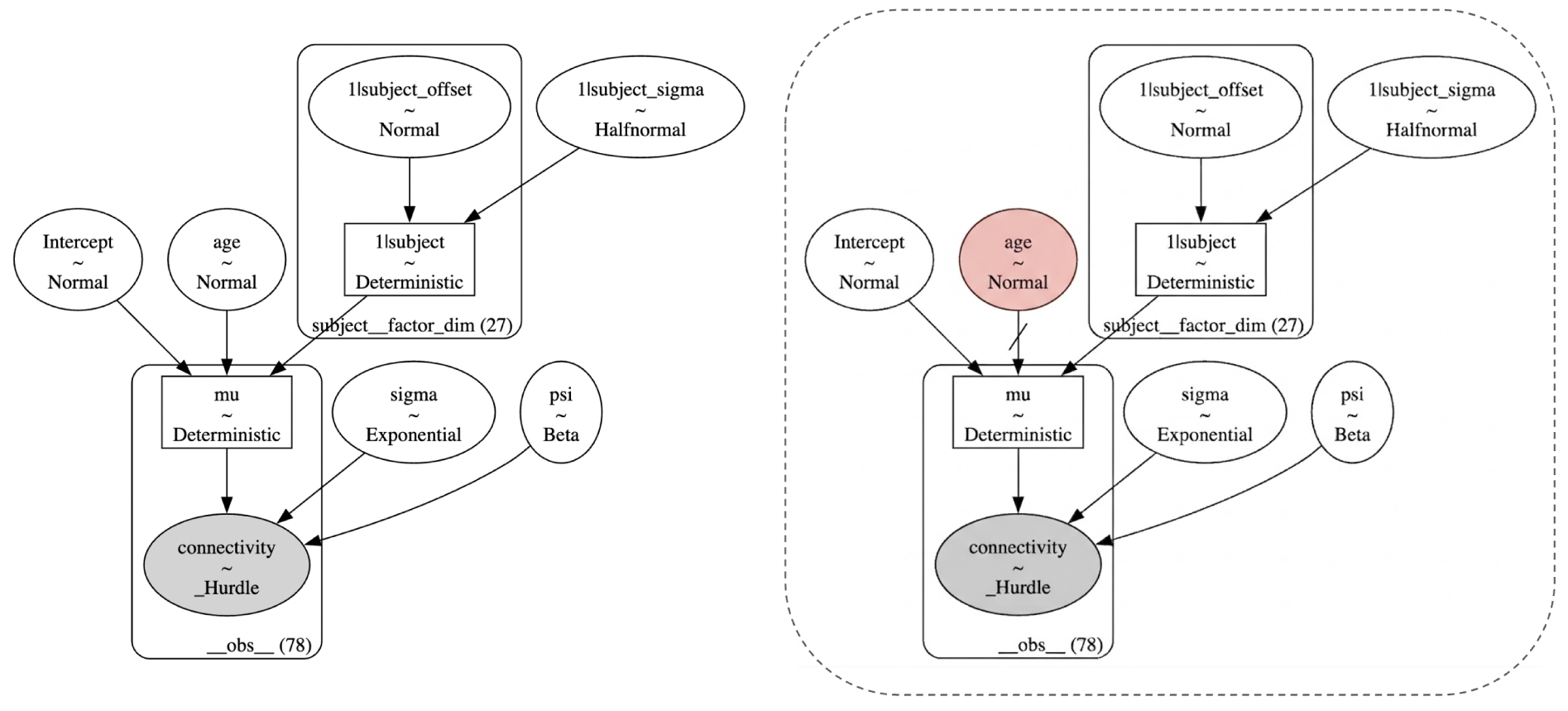
Full model (left) and counterfactual null model (right) graphical representations. The hierarchical structure of each mixed effects model is captured by the directed flowgraph, where (hyper)parameter dependencies are indicated by arrows. An intervention (here shown in red) entails surgically modifying the coefficient values (boxes/circles) and severing dependencies (arrows). In the models above, the null intervention on age only demands fixing the age variable to zero, which is equivalent to dropping it from the graph due to the lack of upstream dependencies. Importantly, interventions do not alter the variance structures of the rest of the model.

Where *β* is a vector of fixed-effect coefficients, *u* captures the random-effect covariance structures, and *X*_obs_ represents the observed design matrix. To obtain a counterfactual generative model, we perform a *do*-operation that entails substituting the observed design matrix *X*_obs_ with the counterfactual design matrix *X*_null_ to alter interactions with the fixed effect of interest while preserving all other covariance structures.

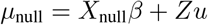

This model is quite simple, so the intervention need only modify the single coefficient of interest, but *do*-operations provide a framework for counterfactual model construction from general hierarchical model structures.

We constructed a counterfactual model from each link’s model by intervening on the covariate of interest. We simulated *K* = 16000 plausible artificial data sets under the null by drawing from the posterior distribution of the log mean *µ*_null_ of each link in the counterfactual model. The log mean captures model-predicted multiplicative effects of age or condition on the observed connectivity in each cell (*i, j*). The result was 16000 null connectivity maps simulated from the counterfactual model. These null maps were used to significance threshold the log mean connectivity from the observed model, *µ*_obs_.

We evaluate each network link (*i, j*) by defining a contrast map which is compared to null contrast maps to set a significance threshold. Here, a contrast is the difference between two groups or conditions, *A* and *B*:

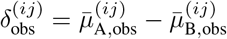

where the empirical posterior expectations for the two levels are calculated from Monte-Carlo draws:

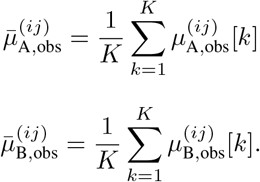

The variance of the post-intervention distribution of *A* can be quantified by comparing each draw 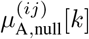 to the static baseline of the observed contrast map, 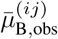:

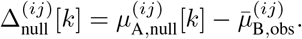

By keeping the observed baseline fixed, the variance of 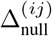 acts as a non-parametric standard error that preserves the covariance structure of the network. The Appendix provides a formal derivation demonstrating how this empirical comparison asymptotically maps to a classical, Wald-type hypothesis test.

#### 2.6.4 Whole-Brain: Cluster Enhancement and Significance Testing

A näıve comparison of 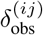 to the distribution 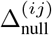 would encounter two primary issues. The most obvious is the need to correct for comparisons across all (*i, j*) pairs, which would likely nullify any initially significant results. The second aspect is related to the first: each regression captures an effect for a single link that is likely individually weak. However, it is conceptually unlikely that distributed changes in the brain should manifest with such narrow scope. Interpreting the single-link regression contrasts *δ*^(*ij*)^s, even with adequate false discovery correction, ignores the broader graph topology that may underlie these changes.

Threshold-Free Cluster Enhancement (TFCE, Smith and Nichols (2009)) is a technique commonly used in fMRI analysis to address such limitations. In fMRI analyses, linear models are fit to each voxel, and effects are spatially integrated across voxels *post-hoc*. TFCE is a popular nonlinear integration method that aims to reweight spatially broad but locally weak effects such that they can be compared to concentrated and strong effects.

Zalesky et al. (2010) offer a framework for clustering *in the graph domain*. For a given link, its graph cluster is defined as the set of links in its strongly connected component and is determined via a breadth-first traversal. The number of links within the link’s strongly connected component defines its network-based statistic (NBS). Baggio et al. (2018) replaced the spatial cluster extent in the traditional TFCE framework with the graph cluster extent given by NBS, which is denoted nbs-TFCE.

Here, nbs-TFCE was applied to the observed data contrast *δ*_obs_ and to each of the null statistic maps Δ_null_[*k*] using the recommended values for graphs of *E* = 0.75 and *H* = 3.25 (from Baggio et al., 2018). The observed nbs-TFCE values were evaluated against the maximum nbs-TFCE value in each: only effects exceeding the 95th percentile of 16000 null maximum values (*α* = 0.05) were considered significant. Figure 2 illustrates this procedure for a single contrast. Coefficient magnitudes, as opposed to signed values, were enhanced since only considering networks comprising positively or negatively signed links in isolation causes fragmentation (e.g., when a negative link connects two positive hubs), thereby underestimating true network extents. Enhanced values were re-signed after significance testing.

**Figure 2:**
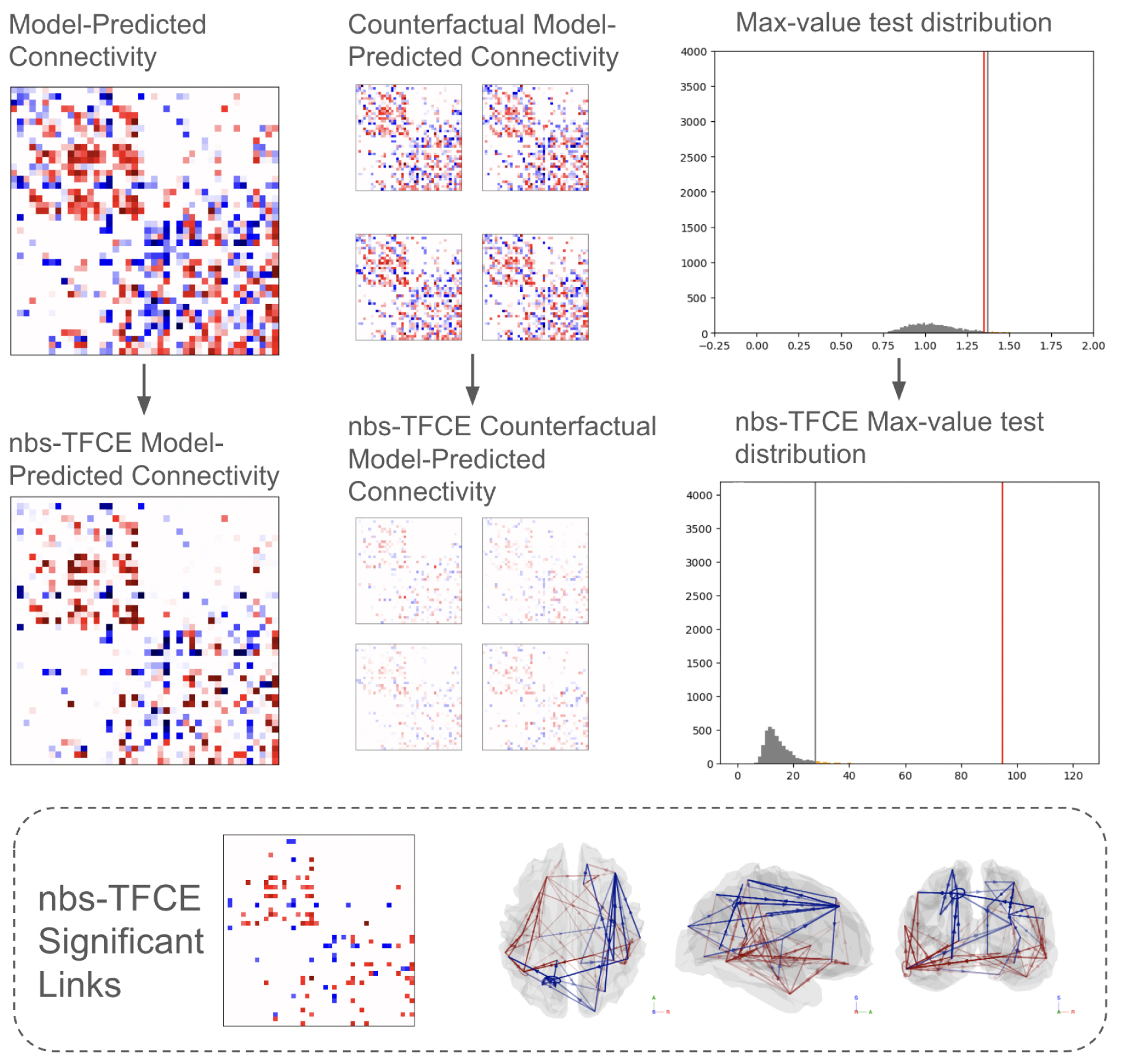
Graphical depiction of the statistical testing procedure used to identify significant cortical networks. **Top Row:** Cbserved and counterfactual (null) model-predicted connectivity contrasts. The histogram to the right compares the unsigned maximum link value in the observed contrast map to the unsigned maximum link values across all null models, producing a maximum that does not pass the *α* = 0.05 significance threshold. **Middle Row:** nbs-TFCE is applied across links (across cells) to enhance network topologies with consistent signal values. Because the observed contrast map contains increased structure relative to the null maps, nbs-TFCE results in comparatively more enhancement to the observed map, as shown in the rightmost histogram. **Bottom Row:** The networks that pass the significance threshold at *α* = 0.05 after nbs-TFCE are shown in matrix form and plotted within a glass brain. Links are re-signed after testing.

### 2.7 NLGC for Stimulus-Driven Response Analysis (Time-Locking)

The Temporal Response Function (TRF) framework uses linear models to estimate the impulse response of the neural system – that is, how the time courses of neural responses temporally track features of a stimulus. In this study, TRFs were fit to the neural source estimates obtained from NLGC to investigate how auditory responses time-locked to features of continuous speech.

#### 2.7.1 TRF Predictors

TRFs were fit to two kinds of predictors designed to capture acoustically-driven auditory responses: the acoustic envelope and envelope onsets.

The acoustic envelope predictor captures amplitude modulation of the speech signal and reflects its time-varying acoustic energy. In contrast, the acoustic envelope onset predictor indexes transients in the speech signal, which are especially prominent at the beginnings of syllables and phonemes.

Both predictors were derived using a gammatone filterbank (Gammatone Filterbank Toolkit 1.0; Heeris (2018)). The speech signal was filtered into 256 frequency channels with logarithmically spaced center frequencies ranging from 20 to 5000 Hz. For each channel, the envelope was extracted, resampled to 1000 Hz, and log-transformed. The resulting 256-band envelope spectrogram was then averaged into 8 logarithmically spaced bands. This yielded the final acoustic envelope predictor.

The acoustic onset predictor was computed from the same 256-band envelope spectrogram by applying an auditory edge detection algorithm (Fishbach et al., 2001; Brodbeck et al., 2023). The resulting onset spectrogram was similarly averaged into 8 logarithmically spaced bands, matching the acoustic envelope predictor.

The distributions of both predictors were verified to be comparable across experimental speech conditions.

#### 2.7.2 TRF Boosting

Temporal response functions (TRFs) are estimated from the data using an iterative boosting algorithm implemented in the Eelbrain python package (Brodbeck et al., 2023). The general procedure is to place a basis function, in our case a hamming window with fixed main lobe width, at a latency which minimizes the error between estimated and true neural responses. This placement strategy is iterated until additional windows result in no decrease in testing error, resulting in a filter comprising many superimposed windows across several time lags. When properly fit, the windows cluster at particular time delays, producing distinct peaks that reflect positive and negative neural potentials occurring at characteristic latencies relative to the stimulus.

For each cortical patch, we retained four eigenmodes, each with a vector spatial orientation. We project these eigenmodes, which have orientations particular to each subject’s neuroanatomy and head orientation, onto canonical RAS (right, anterior, superior) 3-axis unit vectors that are standard across all subjects. This registration facilitates inter-subject comparison and averaging of TRFs.

Following RAS registration, TRFs were estimated using boosting with an *L*_2_ penalty and were fit using a large integration window of −1000 to 1000 milliseconds and Hamming window main-lobe widths of 75 ms. Input and output data were standardized (*Z*-scored) prior to fitting the TRF models.

#### 2.7.3 TRF Significance Testing

TRFs were fit to each of the 84 cortical patches. Significant responses were determined by comparison of the observed model to three ‘null’ TRF models obtained by circularly shifting the predictor time series in increments of one-fourth its total duration, each time fitting a new TRF model. The purpose of this shifting operation is to preserve the statistical properties of the predictor time series but to remove its temporal association with the measured neural responses.

Significant TRF prediction accuracies were determined by comparing the observed model prediction accuracies to those of the averaged null models. To effect this comparison, individual accuracy values were first *z*-transformed and subjected to an related-measures *t*-test. To control for multiple comparisons, we applied TFCE to the resulting *t*-values. Unlike the aforementioned graph applications of TFCE, it was applied here only over spatially contiguous samples. Significance was determined via a max-TFCE statistic permutation test (10,000 iterations) in which condition labels were randomly flipped across subjects. Only clusters exceeding the 95th percentile of the maximum TFCE distribution were considered significant (*α* = 0.05). One test was performed for each band.

TRF prediction accuracies were used to assess whether GC link patterns were confounded by the presence of a shared exogenous input; speech listening drives time-locked neural activity at various nodes over the cortex, and slight variations in the latencies of these stimulus-driven responses could produce illusory GC links. We isolated nodes with significant TRF prediction accuracies and compared connectivity within this significant set *S* to connectivity involving nodes in the non-significant set *N* as the link origin, target, or both. A Kolmogorov-Smirnov test was performed, for older and younger adults separately, to contrast the confounding *S* → *S* case with the control cases *N* → *S, S* → *N, N* → *N*.

## 3 Results

### 3.1 Speech Listening and Resting State Networks

First, whole-brain speech-listening networks were identified in the Theta and Delta bands by contrasting connectivity densities between speech-in-quiet listening and resting-state conditions, where positive differences are identified with the speech-listening networks and negative differences are identified with the resting-state networks (note that, in this formulation, connectivity common to both states do not contribute to either network). The model-derived networks are shown for each band in Figure 3. The plots show the model-predicted average differences in log connectivity density between speech listening and resting state, masked by significant effects identified via nbs-TFCE on model-predicted values (*p* < 0.05 corrected). Because the differences are on a logarithmic scale, darker values indicate multiplicative relative differences in connectivity.

**Figure 3:**
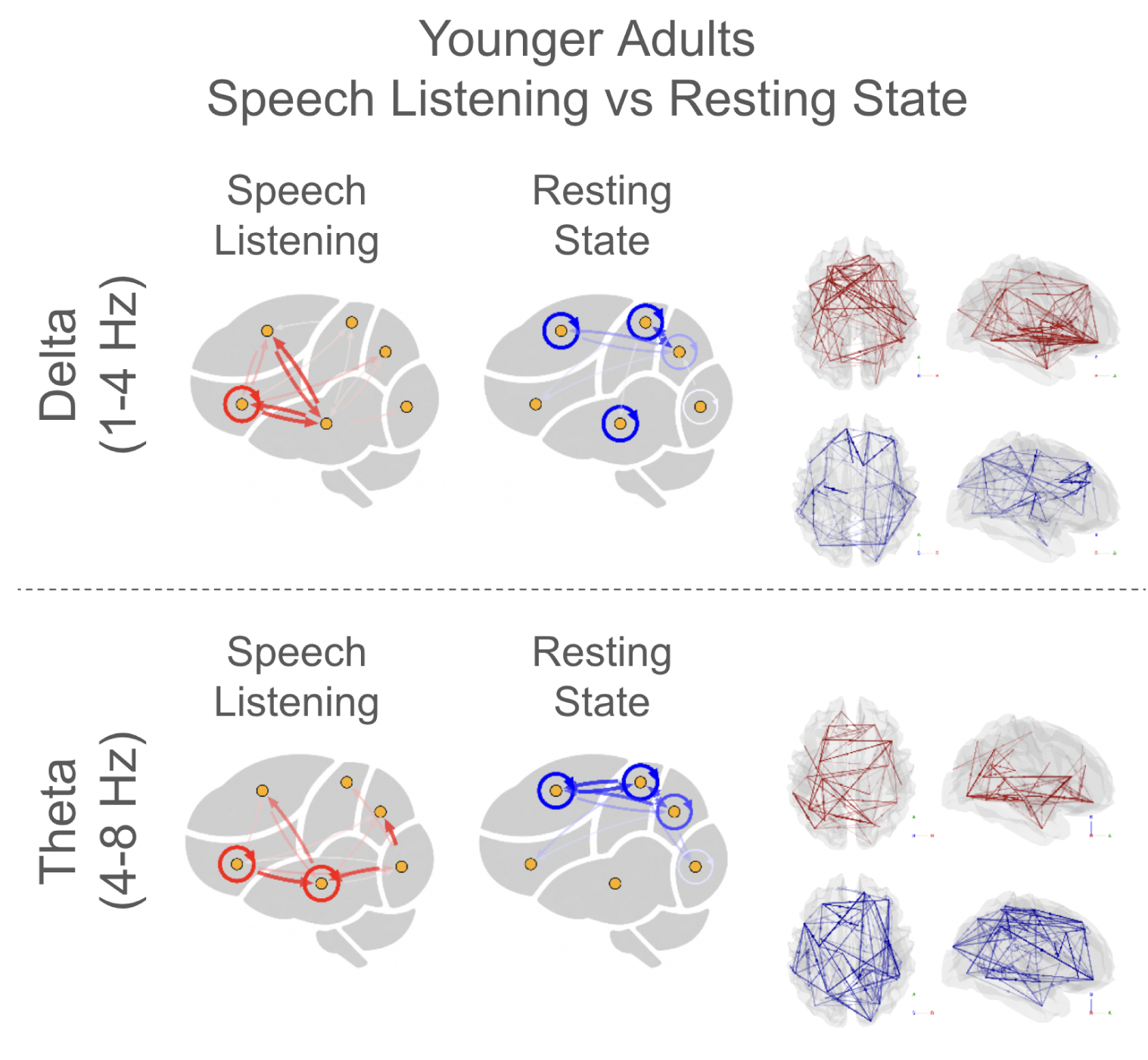
Whole-brain connectivity differences between speech-in-quiet listening and resting state in younger adults across Delta (left panel) and Theta (right panel) frequency bands. Each panel shows model-predicted differences in log connectivity density, masked by significant effects identified using nbs-TFCE (*p* < 0.05, corrected). Warmer colors (red) indicate stronger connectivity during listening, while cooler colors (blue) indicate stronger connectivity in resting state. **Delta band:** Speech listening is associated with bilateral temporal-frontal connectivity. Resting state connectivity is observed in default-mode regions including posterior cingulate cortex, medial prefrontal cortex, and angular gyrus. **Theta band:** In contrast to the Delta band, Theta band speech-listening networks show left-lateralized temporal-frontal and temporal-parietal connectivity. Resting state connectivity, however, is again observed in bilateral default-mode regions, as in the Delta band.

The Delta band contrasts show significantly increased bilateral temporal-frontal connectivity when listening. Neural responses to speech in the Delta band are known to track words and supra-segmental features of speech such as intonation and rhythm (Brodbeck et al., 2018a), supported by interactions between temporal and inferior frontal cortical regions (Fridriksson et al., 2016). At rest, the Delta band connectivity is consistent with canonical default mode resting state networks that comprise areas of posterior cingulate cortex, medial prefrontal cortex, and angular gyri (Horn et al., 2014; Andrews-Hanna et al., 2014)

The Theta band contrasts show significantly increased temporal-frontal and temporal-parietal connectivity. Many of the connectivity increases are observed to originate from or terminate in the left temporal lobe. Neural responses to speech in the Theta band index syllable onsets and sublexical features such as phonemes (Brodbeck et al., 2018a) and correlate with speech clarity (Etard and Reichenbach, 2019). These processes involve temporal, inferior parietal, and inferior frontal cortical regions during speech perception (Fridriksson et al., 2016). At rest, the Theta band connectivity is similarly consistent with canonical default mode resting state networks.

Overall, when listening Delta-band connectivity increases were predominantly bilateral and restricted to temporal-frontal interactions, whereas Theta-band increases were more left-lateralized and additionally involved temporal-parietal connections.

### 3.2 Whole-Brain Speech Listening Networks in Younger Adults

Significant (*p* < 0.05) nbs-TFCE model-derived contrasts between listening conditions in each frequency band, for younger adults, are shown in Figure 4.

**Figure 4:**
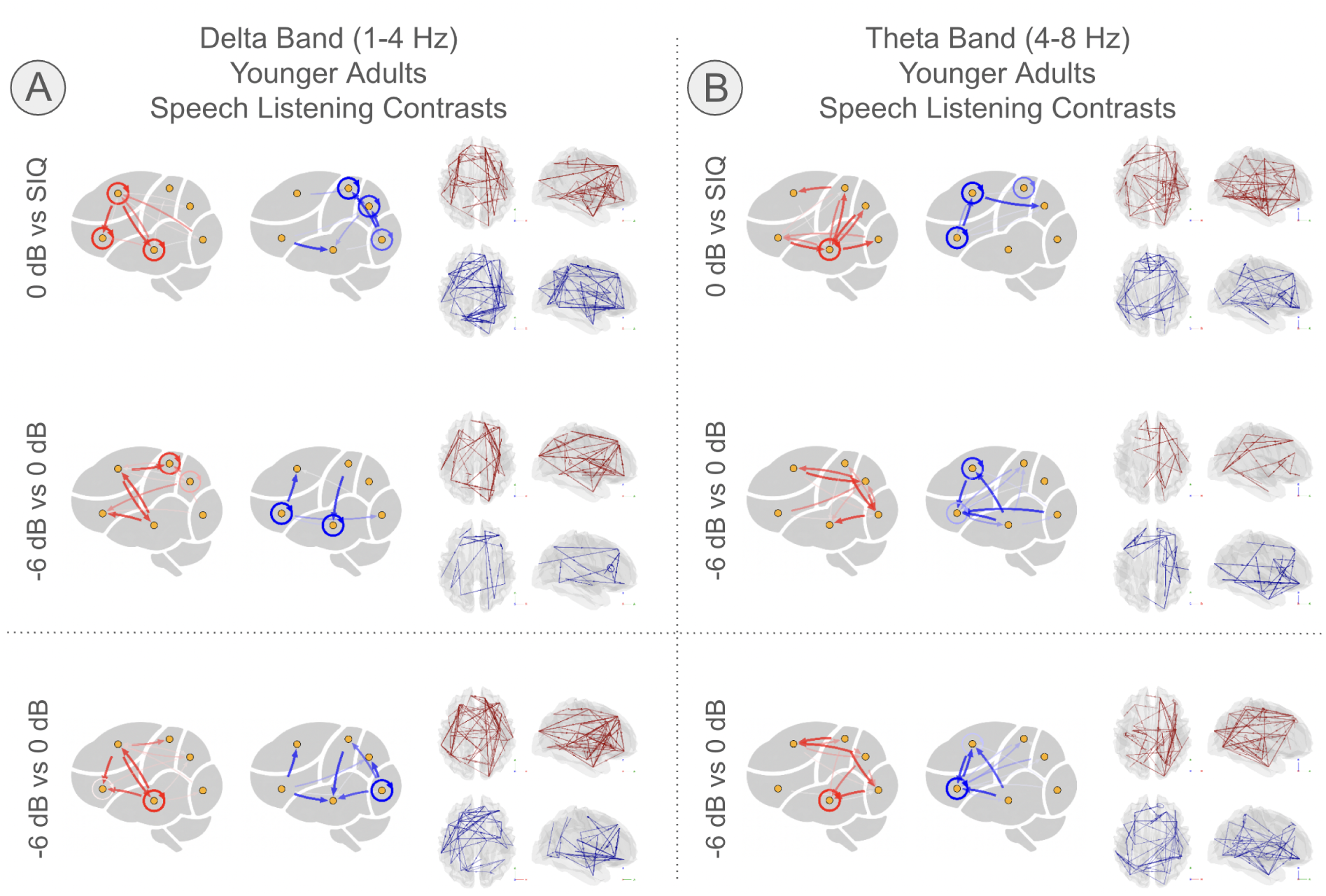
Significant connectivity contrasts between listening conditions in younger adults for (A) Delta and (B) Theta frequency bands. Networks represent model-predicted differences in log connectivity density between conditions, thresholded using nbs-TFCE (*p* < 0.05, corrected). Red edges indicate stronger connectivity in the more difficult listening condition, whereas blue edges indicate stronger connectivity in the easier condition. Rows show condition contrasts (speech-in-quiet vs. 0 dB cocktail, 0 dB vs. −6 dB cocktail, and −6 dB cocktail vs. speech-in-quiet), and columns correspond to frequency bands. **Delta band:** The SIQ vs. 0 dB contrast shows increased bilateral temporal-frontal connectivity involving superior frontal regions. The 0 dB vs. −6 dB contrast shows additional increases in frontal and parietal connectivity with greater interhemispheric involvement. The −6 dB vs. SIQ contrast shows widespread increases centered on superior frontal regions under more difficult listening conditions. **Theta band:** The SIQ vs. 0 dB contrast shows predominantly right-lateralized increases in parietal-temporal and parietal-frontal connectivity. The 0 dB vs. −6 dB contrast shows reductions in bilateral temporal-frontal connectivity with additional right-lateralized parietal involvement. The −6 dB vs. SIQ contrast shows a shift toward right-lateralized connectivity patterns with a precentral hub.

In the Delta band, with increasing listening difficulty, we observe bilateral increases in superior frontal to temporal connectivity and frontal-parietal connectivity. These effects involve superior frontal regions near the junction of the precentral and superior frontal sulci (the frontal eye fields), as well as more anterior areas near the dorsolateral prefrontal cortex (dLPFC), as described next.

Comparing speech-in-quiet (SIQ) listening to the more difficult 0 dB cocktail listening condition (Figure 4; row 1, column A), we observe bilateral increases (red) in connectivity to and from superior frontal areas near the frontal eye fields. In the right hemisphere, these are predominantly connections originating from temporal and parietal areas. These, in turn, connect to left superior temporal and inferior frontal regions that subsequently connect to areas in the left and right temporal lobes. Reduced interhemispheric connectivity is also observed relative to the SIQ condition.

As listening conditions degrade further from 0 dB to −6 dB (Figure 4; row 2, column A), further increases in frontal and parietal connectivity are observed in conjunction with increased interhemispheric connectivity. Increased bidirectionality of connectivity involving the right frontal eye fields is observed. Decreases in connectivity present in anterior cingulate cortex and inferior frontal areas.

The final row presents the bulk effects of speech listening difficulty (row 3, column A), contrasting the −6 dB cocktail condition with the SIQ condition. The significant effects show a large increase in connectivity centered around the frontal eye fields with adverse listening conditions. In easier conditions, increased left temporal to left inferior frontal connectivity is observed, consistent with ventral stream engagement.

In Theta, as listening conditions transition from easier (SIQ) to more difficult (0 dB cocktail) we observe rightward increases in connectivity (Figure 4; row 1, column B) mainly involving areas of right parieto-frontal (dorsal and anterior lateral frontal lobe, precentral sulcus) and right temporo-frontal (inferior and ventral frontal lobe, posterior temporal lobe) circuits (Labache et al., 2024). Connectivity decreases are primarily centered around anterior cingulate and inferior frontal regions.

As listening conditions transition from 0 dB to −6 dB (row 2, column B), some additional marginal increases in right-lateralized parietal-centric connectivity are observed, but, overall, connectivity significantly decreases in bilateral temporal and inferior frontal areas.

The bulk effects of speech listening difficulty (row 3, column B) show increased connectivity in temporal and inferior frontal connectivity in SIQ listening that transitions to rightward patterns of connectivity with a precentral hub of connections, consistent with dominantly dorsal stream engagement. Table 2 presents a summary of selected connectivity changes across band and listening condition.

**Table 1:**
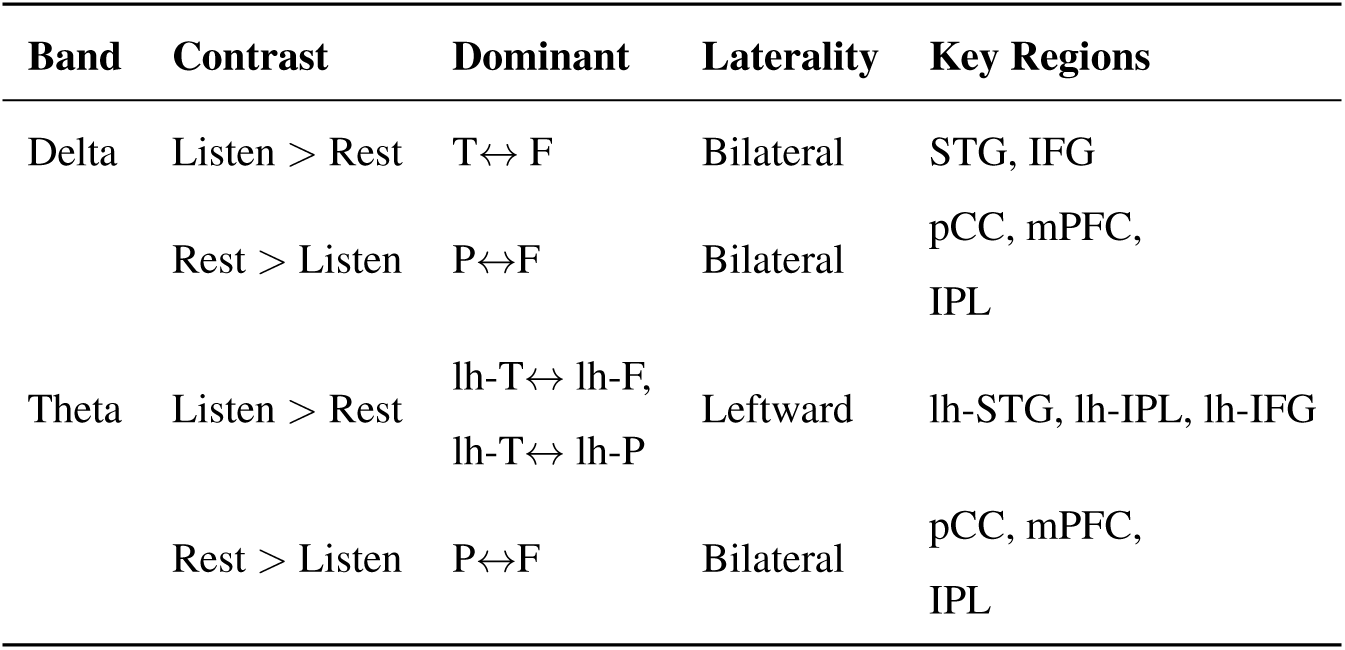
Summary of significant connectivity differences between speech listening and resting state across frequency bands.

**Table 2:** Summary of younger adults’ selected connectivity changes across speech listening conditions by frequency band.

| Band | Contrast | Dominant Changes | Laterality | Key Regions |
| --- | --- | --- | --- | --- |
| Delta | 0 dB > SIQ | T → F, P → F | Bilateral<br>(right-dominant) | STG, FEF, SPL |
|  | -6 dB > 0 dB | F and P increases | Bilateral | FEF, SPL |
|  | -6 dB > SIQ | F-centered increases | Bilateral | FEF |
| Theta | 0 dB > SIQ | P → T, P → F | Right | STG, precentral,<br>IFG |
| | -6 dB > 0 dB | T↔F decreases | Bilateral | STG, IFG |
|  | -6 dB > SIQ | P-centered increases | Right | STG, precentral,<br>IFG |

### 3.3 Whole-Brain Networks Age Comparison

Significant (*p* < 0.05) nbs-TFCE model-derived contrasts between older compared to younger adults for each listening condition (SIQ, 0 dB, and −6 dB) and frequency band are shown in Figure 5, with (Older > Younger) in red and (Younger > Older) in blue. Differences are shown on a logarithmic scale, such that stronger colors reflect multiplicative differences in connectivity.

**Figure 5:**
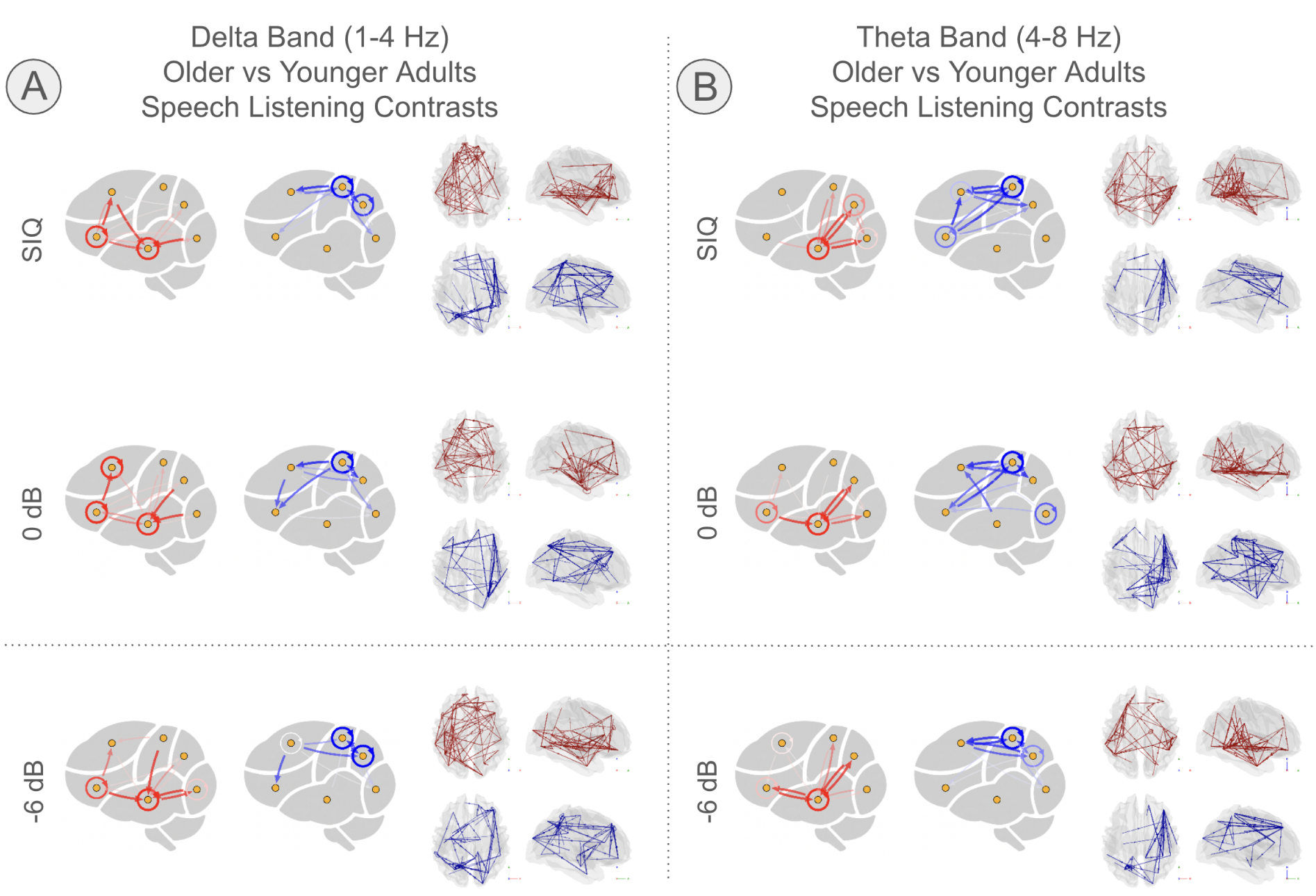
Whole-brain connectivity differences between older and younger adults across listening conditions in the (A) Delta and (B) Theta frequency bands. Each panel shows model-predicted differences in log connectivity density (Older > Younger in red; Younger > Older in blue), masked by significant effects identified using nbs-TFCE (*p* < 0.05, corrected). **Delta band:** Across conditions, older adults show increased connectivity in left temporal-frontal networks, including inferior frontal and insular regions interacting with superior and middle temporal cortices. Younger adults show relatively stronger right-lateralized frontoparietal connectivity. **Theta band:** Older adults show increased local temporal and temporal-parietal connectivity, particularly involving superior and middle temporal regions and inferior parietal areas. Younger adults again show stronger rightward frontoparietal connectivity, with similar spatial organization to that observed in the Delta band. **Across conditions:** Age-related differences exhibit broadly similar spatial patterns within each frequency band. While the extent of significant connectivity varies across listening conditions, the overall organization of age-related network differences remains consistent.

In the Delta band, older adults show consistently increased connectivity in left temporal and frontal areas relative to younger adults across listening conditions, consistent with increased ventral stream engagement. In particular, these regions include the left insula and inferior frontal regions within and near the opercula which interact with left superior and middle temporal areas. In contrast, younger adults, relative to older adults, show increased rightward frontoparietal connectivity including regions near the frontal eye fields, precentral gyrus, and superior parietal lobule. These patterns resemble the band-specific connectivity profiles observed in younger adults’ speech listening versus rest contrasts.

In the Theta band, local temporal and temporal-parietal connectivity is increased in older adults relative to younger adults. These regions include superior and middle temporal areas that connect to the inferior parietal lobules and frontoparietal areas, consistent with increased dorsal stream engagement A similar rightward frontoparietal network shows decreased connectivity in older adults relative to younger adults, with similar morphology to the network identified in the Delta band.

Across listening conditions, age-related differences exhibit broadly similar spatial patterns across Delta and Theta bands, with consistent differences in network topology between older and younger adults observed within each condition. While the extent and strength of these differences varied across conditions, we observe no clear qualitative shift in the organization of age-related connectivity patterns.

Overall, age-related differences are characterized by increased temporal and temporal-frontal connectivity in older adults and increased right-lateralized frontoparietal connectivity in younger adults, with this contrast preserved across listening conditions. Table 3 presents a summary of the changes observed between younger and older listeners in both frequency bands.

**Table 3:**
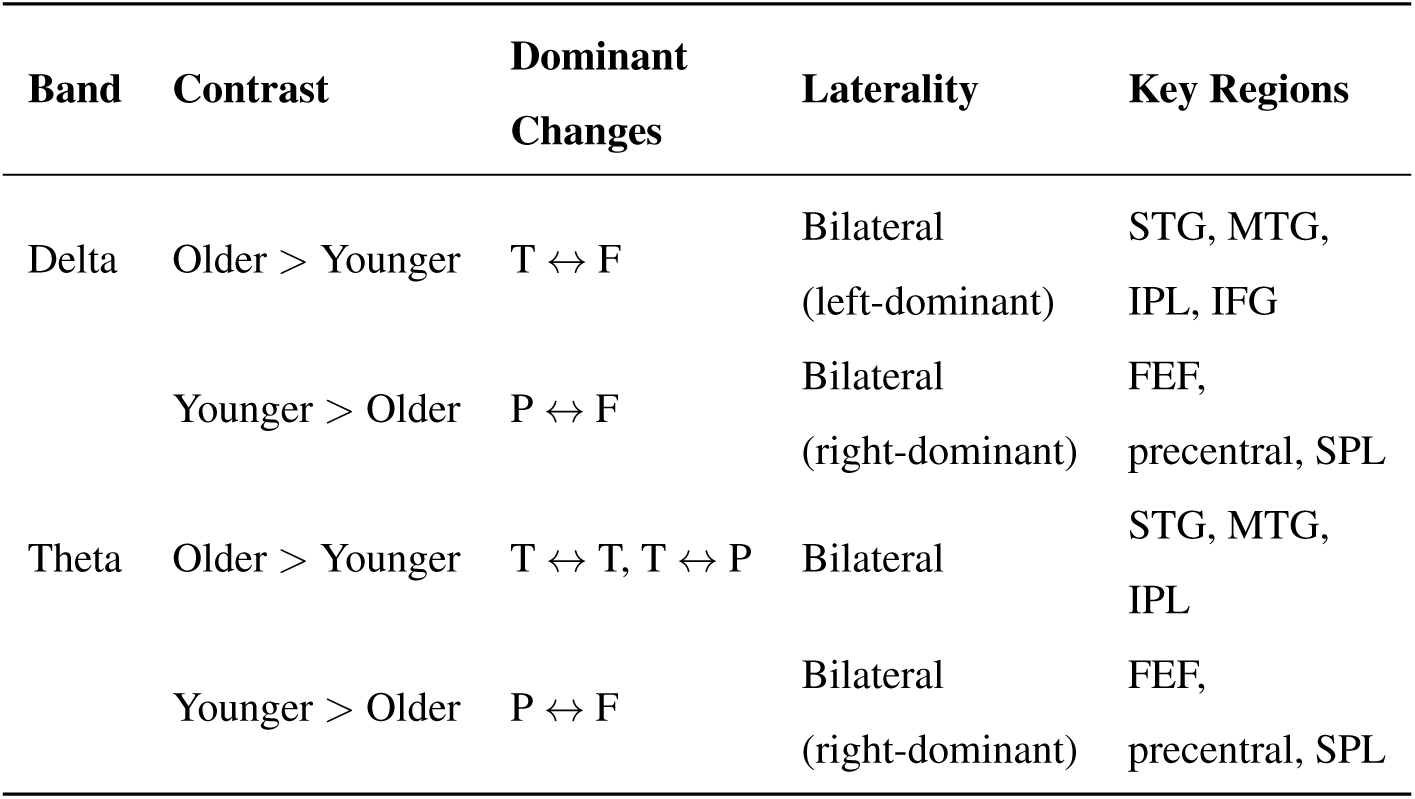
Summary of selected connectivity changes between older and younger listeners by frequency band.

| Band | Contrast | Dominant Changes | Laterality | Key Regions |
| --- | --- | --- | --- | --- |
| Delta | Older > Younger | T ↔ F | Bilateral<br>(left-dominant) | STG, MTG,<br>IPL, IFG |
|  | Younger > Older | P ↔ F | Bilateral<br>(right-dominant) | FEF,<br>precentral, SPL |
| Theta | Older > Younger | T ↔ T, T ↔ P | Bilateral | STG, MTG,<br>IPL |
|  | Younger > Older | P ↔ F | Bilateral<br>(right-dominant) | FEF,<br>precentral, SPL |

### 3.4 Time-Locking Analysis

Time-locked cortical speech tracking measures were then found for the sources and targets of the significant directed connectivity links. Figure 6, left presents cortical patches where grand-average TRF prediction accuracies (*r* values) were significant, with average *r* values plotted for Delta and Theta band TRFs. Significance was determined via max-TFCE cluster test over 10,000 permutations, with Theta max *p* < 0.0001 and Delta max *p* = 0.0001. Responses are maximally predicted in areas in the vicinity of the superior temporal lobe.

**Figure 6:**
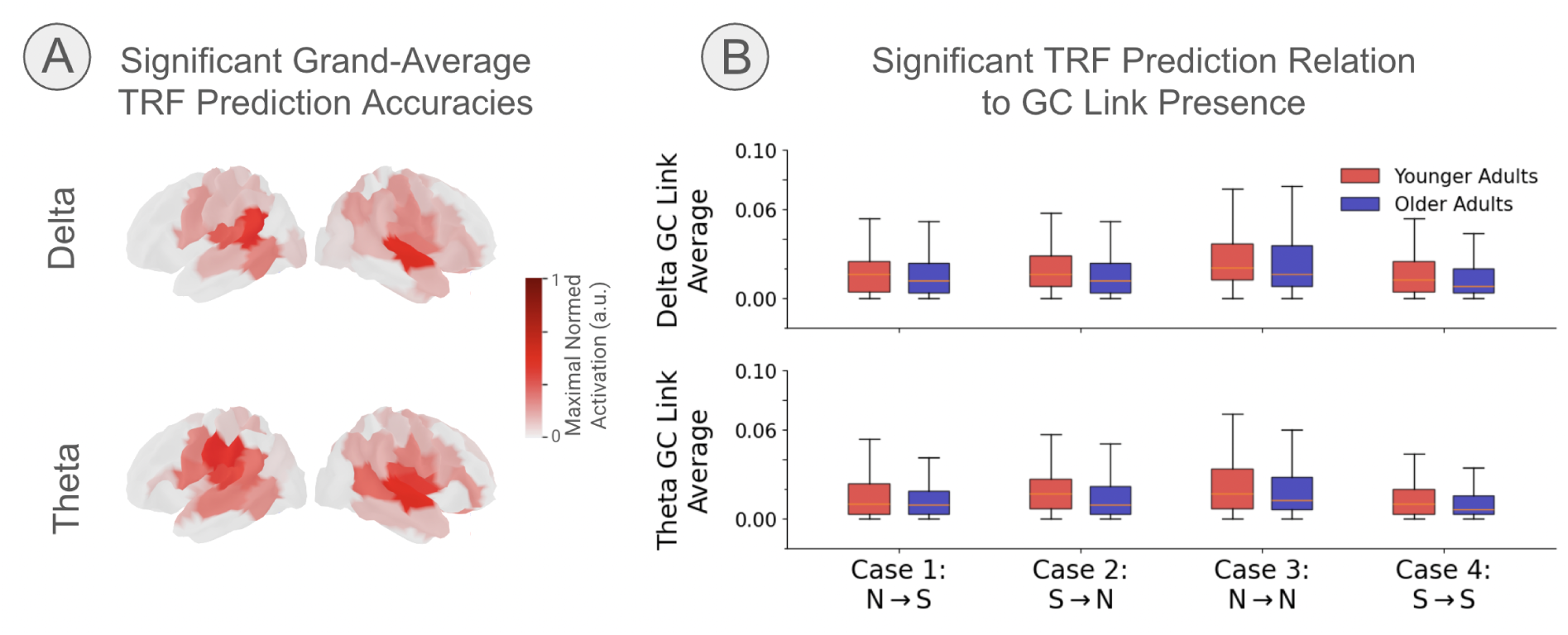
Time-locked cortical speech tracking in Delta and Theta bands, related to significant directed connectivity links. (A) Cortical regions where grand-average temporal response function (TRF) prediction accuracies (*r* values) were significant, with mean Delta- and Theta-band prediction accuracy masked by significance. Significant responses were centered near superior temporal cortex in both bands. (B) Spatial overlap between significant NLGC links and temporal-lobe regions with significant (*S*) and non-significant (*N*) TRF prediction accuracy. Four cases are shown depending on whether the neither source nor target node, source node, target node, or both nodes overlapped with significant TRF responses. Case 4 (*S* → *S*) represents a potential confound in which both nodes are driven by the same auditory stimulus. No significant incidence of such links was observed within either age group in either band (Delta older adults, KS *p* = 0.906; Delta younger adults, KS *p* = 0.990; Theta older adults, KS *p* = 0.995; Theta younger adults, KS *p* = 0.994), suggesting that the connectivity results were not driven solely by shared sensory responses.

The overall increased neural tracking fidelity in older adults, exemplified by higher prediction accuracy and response magnitude in their TRF models (see Supplement), is a well-known finding thought to arise, at least in part, from a central gain mechanism.

It is possible that connectivity between cortical areas is being driven by an exogenous stimulus unknown to the Granger-Causal model. In this case, the stimulus could drive neural responses in two cortical areas with a small delay, resulting in spurious connectivity between the two regions. We measure the fidelity of the stimulus representation in cortex using TRFs, and count the number of links with source and target involving nodes in the set of nodes *S* with significant stimulus-driven responses. We contrast these links with links with at least one source or target in the set *N* of non-significant stimulus-driven responses. The results are shown in the right panel of Figure 6. Case 4 illustrates the confounding case where both source and target of an NLGC link are being driven by an exogenous auditory stimulus (*S* → *S*). Cases 1 through 3, in contrast, are not confounded. Because older adults have enhanced cortical envelope representations relative to younger adults, we compare the incidence of these connections for both age groups in each band. No significant incidence of confounding links was observed within either age group in either band (Delta older adults, KS *p* = 0.906; Delta younger adults, KS *p* = 0.990; Theta older adults, KS *p* = 0.995; Theta younger adults, KS *p* = 0.994) suggesting that the connectivity results were not driven solely by shared sensory responses.

## 4 Discussion

Distinct condition-dependent changes in connectivity are observed in younger whole-brain speech-listening networks compared to resting state, and also when difficulty transitions from easy-to-moderate and moderate-to-hard. Furthermore, older adults exhibit distributed changes compared to younger adults at each difficulty test point.

### 4.1 Younger Adults’ Speech Listening

In the Delta band, speech-listening networks comprise connections between bilateral inferior frontal and temporal regions. These are areas primarily associated with the ventral auditory processing stream of the dual-stream auditory processing model (Rauschecker and Tian, 2000; Arnott et al., 2004; Rauschecker and Scott, 2009). Neural activity along the ventral stream is thought to mediate semantic and lexical representations of speech (Fridriksson et al., 2016; Wang et al., 2024). Delta-band neural activity has been observed to time-lock to word onsets and other supra-segmental features of speech such as intonation and rhythm (Brodbeck et al., 2018a), and so temporofrontal connectivity in the Delta band may undergird the semantic processing of speech. Despite this time-locking-based motivation, however, we also demonstrated that most of the significant connections observed do not show time-locking in the connection’s source or target, or show time-locking in only one but not the other, so the frequency band itself may be more important than what time-locked responses in that frequency band show..

The lack of lateralization we observe in Delta speech-listening networks may be due to differences between the topology of networks that support processing of speech-relevant features, which may lack strong lateralization, and the locus of time-locked word processing that is known to be strongly lateralized. For instance, recent studies have shown that bilateral inferior frontal regions time-lock to other low-frequency features of speech, such as chunks composed of several words, independent of prosodic cues or acoustics (Chalas et al., 2024), in contrast to the strong left-lateralization of word-processing cortical areas (e.g., Peelle et al., 2009; Peelle, 2012; Karunathilake et al., 2024).

Speech-listening networks in the Theta band exhibit left lateralization, with most connections being left temporoparietal or temporofrontal. Temporal-to-parietal connections in particular are associated with the dorsal auditory processing stream (STG, IPL, PMC) and are thought to relate to phonological (language representations pertaining to phonemes) processing of speech (Fridriksson et al., 2016). Neural responses in the Theta band have been shown to time-lock to sublexical features of speech such as phonemes and syllable onsets (Brodbeck et al., 2018a) and correlate with speech clarity (Etard and Reichenbach, 2019). In this way, Theta-band connectivity is intrinsically linked to rhythmic processing and may reflect speech processing within the dorsal stream. Again, however, most of the significant connections observed do not show time-locking in the connection’s source or target, or show time-locking in only one but not the other, so the frequency band itself may be more important than what time-locked responses in that frequency band show.

### 4.2 Younger Adults’ Resting-State Networks

At rest, the Delta and Theta band networks show spatial patterns broadly consistent with canonical fMRI-measured default mode resting state networks that comprise areas of posterior cingulate cortex, medial prefrontal cortex, and angular gyri (Horn et al., 2014). There is some supporting evidence from simultaneous fMRI-EEG studies that the default-mode network morphology presents similarly among Delta (Neuner et al., 2014) and Theta bands (White et al., 2012). It need not have been the case that the Delta and Theta band networks be consistent with each other, let alone with the analogous fMRI results, but this does indeed appear to be so.

### 4.3 Younger Adults’ Speech-Listening Network Connectivity Changes Under Adverse Listening Conditions

Neural connectivity in younger adults is characterized by network-organizational shifts in response to changes in listening difficulty across a wide dynamic range. Reconfiguration consists of increased integration between temporal auditory regions and dorsal frontal and parietal control areas.

In the Delta band, decreasing SNR in attended speech is associated with increased connectivity between temporal regions and hubs in bilateral superior frontal and dorsolateral prefrontal cortex. These areas include the frontal eye fields which form primary component of the dorsal attention network (Petersen and Posner, 2012; Szczepanski et al., 2013). As listening becomes more difficult, we observe increases in connectivity between temporal areas and regions near bilateral FEF (in and around the precentral sulcus and superior frontal sulcus, which in our data are coarsened to include parts of the dLPFC).

The FEF in humans are well-known to be involved in visuospatial attentional orienting and visual perceptual modulation (for review, see Petersen and Posner, 2012; Vernet et al., 2014). There is mounting evidence for auditory spatial attention engaging areas in the FEF absent visual stimulus (Michalka et al., 2015; Lee et al., 2013; Garg et al., 2007; Bharadwaj et al., 2014) and even auditory sensory responses in the FEF Kirchner et al. (2009).

The interactions of temporal areas with bilateral FEF and dLPFC areas may reflect top-down attentional modulation of slower, supra-segmental speech features (e.g., prosody and phrasal structure), which are tracked at Delta timescales (Chalas et al., 2024).

In the Theta band, similar patterns of connectivity are observed, with increases in connectivity between temporal areas and frontoparietal hubs with increasing listening difficulty. Several factors may contribute to the apparent right lateralization of the networks. Lateralization could arise from significance thresholding, noting the presence of left-hemisphere subnetworks in similar areas but with a weaker contrast. However, there is some evidence for a rightward lateralization of frontoparietal attention networks (Labache et al., 2024; Ahveninen et al., 2013).

The apparent lateralization observed relative to the Delta band networks could relate to task- and information-specific modulations in the Theta band. In visuospatial attention, the right hemisphere components of the dorsal attention network (FEF and IPS) are driven stimuli spanning subjects’ entire visual field, whereas homologous areas in the left hemisphere respond to contralateral stimuli (Bartolomeo and Seidel Malkinson, 2019; Sheremata et al., 2018). Similar mechanisms could relate to attention orienting in auditory scenes (Sussman, 2017) and could interact with particular speech features encoded in each frequency band.

The final contrast in both Delta and Theta, performed between speech-in-quiet and −6 dB cocktail listening, reflects network-organizational changes under bulk effects of the most difficult listening conditions tested. In both bands, distinct frontoparietal increases with difficulty are observed, which we relate to uptake in attentional modulation above, but interestingly a decrease in left-hemisphere temporal to inferior frontal connectivity is also present. Connectivity decreases in these areas are related to the auditory ventral stream interpretation above, and therefore may be indicative of poorer semantic processing in the hardest listening condition where the ability to segregate words within attended speech is diminished.

Our findings contribute to a growing body of evidence supporting involvement of frontoparietal areas of the dorsal attention network during auditory tasks as a locus of attention orienting beyond audiospatial or visuospatial modulation.

### 4.4 Network Connectivity Differs Between Older and Younger Adults

Older adults exhibit consistent differences in network topology compared to their younger counterparts across all listening conditions. Their networks emphasize areas in the canonical ventral and dorsal auditory streams, possibly reflecting a combination of top-down connectivity aligning with semantic prediction used to correct degraded speech representations (Park et al., 2015) and bottom-up connectivity analogous to central gain mechanisms that enhance auditory encoding in older adults (Harris et al., 2022).

In the Delta band, older adults exhibit increases in connectivity between primarily left temporal and left inferior frontal areas. As discussed above, these regions are primarily associated with the ventral auditory processing stream of the dual-stream model (Rauschecker and Tian, 2000; Arnott et al., 2004; Rauschecker and Scott, 2009), which is implicated in the mapping of acoustic input onto lexical and semantic representations (Fridriksson et al., 2016; Wang et al., 2024). Correspondingly, Delta band neural activity has been shown to time-lock to word onsets and supra-segmental features of speech such as intonation and rhythm (Brodbeck et al., 2018a). Together, these findings suggest that older adults’ observed connectivity may support the integration of acoustic input with higher-level semantic representations during speech processing. The enhancement of these networks relative to younger subjects could indicate increased top-down semantic prediction whereby speech semantics are used to anticipate or correct portions of the auditory stream, even during speech-in-quiet listening.

In the Theta band, older adults present bilateral networks involving primarily temporal and parietal areas. These regions are associated with the dorsal auditory processing stream, which is thought to perform phonological (phonemic) processing of speech (Fridriksson et al., 2016). Since Theta band responses are known to time-lock to sublexical features of speech such as phonemes and syllable onsets (Brodbeck et al., 2018a) and correlate with speech clarity (Etard and Reichenbach, 2019), it may be that responses in this frequency band are particularly suited to processing within the dorsal stream.

The enhancement of these networks in older adults may be a consequence of diminished fidelity in the speech signal, perhaps arising from structural or functional decline in afferent input to cortex from subcortical areas, or even diminished signal at the periphery (though our older adults had clinically normal hearing, some minor hearing loss is nonetheless observed in our cohort). These changes are the basis for the central gain hypothesis in older adults, which states that age-related peripheral deafferentation contributes to reduced inhibition, resulting in increased cortical response amplitudes (Harris et al., 2022). We observe increases in time-locked neural responses (TRFs) consistent with central gain in older adults across all listening conditions. The locations where time-locked responses were measured did not significantly coincide with either source, target, or both source and target locations of significant links, indicating that the Theta-band connectivity increases in older adults are unlikely to be solely explained by central gain mechanisms. Rather, the presence of these networks within the dorsal auditory stream at all listening difficulties is consistent with increased reliance on phonological processing with aging. The bilaterality of the Theta networks is discussed below.

In younger adults, networks appear relatively stable between Delta and Theta bands. The areas involved are similar to network morphologies observed in attention areas of younger adults’ listening condition network contrasts. We discuss the lateralization of responses next.

### 4.5 Network Connectivity and Aging Models

Several prominent models of cognitive aging provide useful frameworks for interpreting the connectivity differences observed between younger and older adults. HAROLD (Hemispheric Asymmetry Reduction in Older adults; Cabeza, 2002) proposes that age-related changes in measured neural responses arise from compensation, dedifferentiation (the process by which neural representations become less distinctive with aging), or both in combination, resulting in less regional specialization and a reduction in the hemispheric asymmetry of responses. Consistent with this view, older adults in our study showed increased bilateral temporoparietal and temporofrontal connectivity in Theta. These bilateral patterns may reflect a compensatory response that supports performance when sensory input is degraded, or may indicate dedifferentiation-related engagement of homologous functional areas in the contralateral hemisphere.

The CRUNCH model (Compensation-Related Utilization of Neural Circuits Hypothesis) further suggests that older adults recruit additional neural resources at lower task demands than younger adults, but may approach capacity limits sooner as demands increase (Reuter-Lorenz and Cappell, 2008). This perspective aligns with the observed enhancement of older adults’ ventral and dorsal auditory-stream networks even during easier speech-in-quiet listening. Rather than emerging only under the most adverse conditions, cortical recruitment was present across the tested range of listening conditions, suggesting earlier or sustained utilization of compensatory resources.

FUEL (Framework for Understanding Effortful Listening) similarly emphasizes that listening effort reflects an interaction between task demands, sensory fidelity, and available cognitive resources (Pichora-Fuller et al., 2016). In this context, maximal temporofrontal network recruitment was observed in younger adults under moderately difficult listening conditions, compared to the most adverse or easiest conditions, supporting the notion that effort and performance are optimized when task demands remain challenging but still tractable within the listener’s available motivational and cognitive resources. When demands become too great, engagement may no longer increase proportionally, or may decrease, resulting in redistributed network recruitment.

### 4.6 Conclusions

We show that speech listening is associated with reorganization of large-scale networks relative to resting state, with task engagement shifting connectivity away from default-mode-like configurations toward distributed auditory and attentional systems. Throughout our results, speech-listening networks often exhibit frequency-specific topologies. Delta networks involve temporofrontal areas within the auditory ventral stream, coinciding with regions specialized for processing speech features that elicit responses at Delta rates such as words or semantics. Theta networks comprise temporoparietal areas within the auditory dorsal stream, incorporating regions that process speech features that occur at Theta rates, such as phonemes. As listening difficulty increases, these speech-listening networks in younger adults are shown to engage additional dorsal frontoparietal regions, suggestive of top-down attentional modulation of speech processing. Contrasting younger and older listeners reveals stable, broadly-distributed connectivity differences across conditions, with older adults displaying increased connectivity within both ventral and dorsal stream regions that may reflect increased reliance on internal representations of speech to support perception. These age-related differences are consistent with a shift from condition-dependent reconfiguration, exemplified in younger adults, toward an elevated, compensatory baseline engagement of semantic and phonological processes. Within the framework of aging models such as HAROLD, the observed bilateralization and reduced lateral specificity in older adults likely also reflect a combination of compensatory recruitment and dedifferentiation. Together, these findings suggest that age-related changes in speech-listening networks arise from both adaptive and degenerative processes, resulting in altered balance between sensory-driven and top-down contributions to speech perception.

## Acknowledgments

The first version of this project was led by Behrad Solemani, who is greatly missed. This work was supported by the National Institute on Aging (NIA) grant P01-AG055365, the National Institute on Deafness and Other Communication Disorders (NIDCD) grant R01-DC019394 and training grant T32-DC00046 (to VC and CF), and the National Science Foundation (NSF) grant SMA1734892 (to JZS).

## Appendix

We illustrate the connection between our bayesian hypothesis test and similar frequentist approaches by deriving the widely-used Wald statistic from our testing framework.

We are testing H_*a*_ : *θ* ≠ *θ*_0_ and H_0_ : *θ* = *θ*_0_ where *θ* is an unknown parameter of interest and *θ*_0_ is its specified value under the null. For instance, if a participant demographic covariate is hypothesized to exert no effect on network edge connectivity, then *θ*_0_ = 0.

We have *P* (*θ*|*Y*), the posterior distribution. When estimated via Monte-Carlo sampling using *K* draws, the posterior expectation is estimated by the sample mean of the draws.

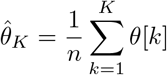

The Bernstein-von Mises theorem links bayesian with frequentist inference by proving that, as the data sample size *N* increases, the posterior uncertainty of a parameter converges to a multivariate normal distribution centered at 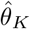 with covariance matrix *N* ^*−*1^ℐ (*θ*_0_)^*−*1^. In our counterfactual generative model, we explicitly enforce the null condition using a *do*-statement, allowing ℐ (*θ*_0_)^*−*1^ to be directly estimated via Monte-Carlo sampling of the counterfactual posterior trace.

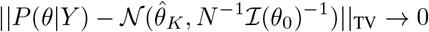

In our counterfactual model we have set the true population parameter, so the Fisher Information can be estimated directly by Monte-Carlo sampling of the posterior for *θ*_0_.

We now show how the Wald statistic can be obtained from trivial manipulations.

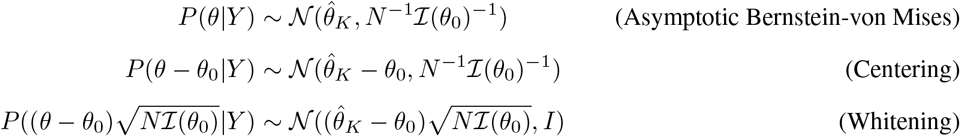

Let 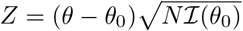 be the whitened residual map. Then:

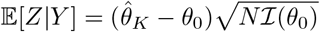

and

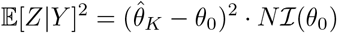

is the Wald statistic. This formalizes how our empirical comparison of a fixed observed point estimate against a counterfactual Monte-Carlo trace respects the underlying information geometry of a frequentist quadratic hypothesis test.

